# A powerful subset-based gene-set analysis method identifies novel associations and improves interpretation in UK Biobank

**DOI:** 10.1101/799791

**Authors:** Diptavo Dutta, Peter VandeHaar, Lars G. Fritsche, Sebastian Zöllner, Michael Boehnke, Laura J. Scott, Seunggeun Lee

**Affiliations:** Center for Statistical Genetics, University of Michigan, MI, USA; Dept. of Biostatistics, University of Michigan, MI, USA; Dept. of Biostatistics, Johns Hopkins University, MD, USA; Graduate School of Data Science, Seoul National University, Seoul, Republic of Korea

## Abstract

Tests of association between a phenotype and a set of genes in a biological pathway can provide insights into the genetic architecture of complex phenotypes beyond those obtained from single variant or single gene association analysis. However, most existing gene set tests have limited power to detect gene set-phenotype association when a small fraction of the genes are associated with the phenotype, and no method exists which identifies the potentially “active” genes that might drive a gene-set-based association. To address these issues, we have developed Gene-set analysis Association Using Sparse Signals (GAUSS), a method for gene-set association analysis that requires only GWAS summary statistics. For each significantly associated gene set, GAUSS identifies the subset of genes that have the maximal evidence of association and can best account for the gene set association. Using pre-computed correlation structure among test statistics from a reference panel, our p-value calculation is substantially faster than other permutation or simulation-based approaches. In simulations with varying proportions of causal genes, we find that GAUSS effectively controls type 1 error rate and has greater power than several existing methods, particularly when a small proportion of genes account for the gene set signal. Using GAUSS, we analyzed UK Biobank GWAS summary statistics for 10,679 gene-sets and 1,403 binary phenotypes. We found that GAUSS is scalable and identified 13,466 phenotype and gene-set association pairs. Within these genes sets, we identify an average of 17.2 (max=405) genes that underlie these gene set associations.

## Introduction

Over the last fifteen years, genome-wide association studies (GWAS) have identified thousands of genetic variants associated with hundreds of complex diseases and phenotype^1^. However, the variants identified to date, individually or collectively, typically account for a small proportion of phenotype heritability^2^. A possible explanation is that due to the large number of genetic polymorphisms examined in GWAS and the massive number of tests conducted, many weak associations are missed after multiple comparison adjustments^3^.

Gene-set analysis (GSA) can identify sets of associated genes that may not be identified using single variant and single gene analysis, especially for rare variants or variants or genes with weak to moderate effects^4^. In GSA, individual genes are aggregated into groups sharing certain biological or functional characteristics. This approach considerably reduces the number of tests performed since the number of gene-sets analyzed are much smaller than the number of genes or genetic variants tested ^5,6^. Additionally, most complex phenotypes are manifested through the combined activity of multiple genes or variants, so that GSA can provide insight into the involvement of specific biological pathways or cellular mechanisms to the phenotype ^7^.

GSA aims to find evidence regarding one of two types of null hypotheses^6^: 1) the competitive null hypothesis in which genes in a gene-set of interest are no more associated with the phenotype than any other genes outside of it; or 2) the self-contained null hypothesis in which none of the genes in a gene-set of interest is associated with the phenotype. Several statistical methods to perform GSA for self-contained null hypothesis have been developed and have successfully identified gene-sets associated with complex diseases^8–15^. For example, de Leeuw et al.^13^ developed MAGMA, a method that transforms p-values of the genes in the gene-set to z-values using an inverse normal transformation and employs linear regression to test the association. Pan et al. ^12^ developed aSPUpath, which uses an adaptive test statistic based on the sum of powered scores and calculates a permutation-based p-value.

However, there are several concerns regarding the power, type-I error control, and computational scalability of these methods. Existing GSA methods often have relatively low power, ^9^ especially in situations where only a few genes within the gene-set have moderate to weak associations with the phenotype ^14^. Additionally, in the presence of correlation between variants or genes due to linkage disequilibrium (LD), many existing methods cannot appropriately control the type-I error^16^. Resampling-based strategies can be used for p-value calculation^17^, but in current implementation these approaches are computationally very expensive, reducing the applicability of the method, especially for large datasets.

Although the existing GSA methods produce a p-value for association between the gene-set and the phenotype, it remains important to identify the specific genes that possibly drive the association signal within the gene-set. This is critical in further downstream analysis and using the results for functional follow-up or suggesting therapeutic targets. Existing GSA methods fail to identify such genes.

Here we describe a computationally efficient subset-based gene-set association method, <u>G</u>ene-set analysis <u>A</u>ssociation <u>U</u>sing <u>S</u>parse <u>S</u>ignals (GAUSS), which aims to increase power over existing methods while maintaining proper type-I error control and facilitate interpretation by extracting a subset of genes that drive the association. GAUSS focuses on the self-contained null hypothesis, as our main goal is to identify phenotype-associated genes or loci. GAUSS identifies a subset of genes (called core subset) within the gene-set which produce the maximum signal of association. The gene-set p-value is calculated through a combination of fast copula-based simulation and statistical approximation approaches using the generalized pareto distribution ^18,19^. GAUSS is constructed using the gene-based test p-values for the genes in the gene-set. The gene-based p-values can be directly computed from the individual level genotype data if available or approximated using GWAS summary statistics (effect sizes, standard errors, and minor allele frequency). Using pre-computed genetic correlation matrices makes GAUSS computationally fast and applicable to large biobank-scale datasets.

Through computer simulation, we show that GAUSS can be more powerful than existing methods while maintaining the correct type-I error. We applied GAUSS to UK-Biobank GWAS summary statistics for 1,403 phenotypes^20^ with 10,679 gene-sets derived from the molecular signature database (MsigDB v6.2) ^21^, demonstrating that GAUSS is feasible for large-scale data and can provide new insights into the genetic architecture of the phenotypes. We have made the association analysis results publicly available through a visual browser.

## Results

### Overview of the methods

To conduct GAUSS, we need p-values for the regions or genes in the gene-set. Popular gene-based tests including genetic association tests like SKAT ^22^, SKAT-Common-Rare ^23^, or genetic expression tests like prediXcan ^24^, can be used to obtain the p-values when individual level data are available. If only GWAS summary statistics (effect size, standard error, p-value, minor allele frequency for each variant) are available, we can approximate the gene-based tests and obtain their p-values using LD information from a suitable reference panel (See Methods) ^25^. The GAUSS test for a given gene-set can be calculated in the following two steps.

#### Step 1, test statistic calculation

To construct the GAUSS test statistic, we start with the gene-based p-values for *m* genes in the gene-set *H* and convert them to z-statistics as *z*_*j*_ = −Φ^−1^(*p*_*j*_) for *j* = 1, 2, …, *m*, where Φ is the standard normal distribution function. The GAUSS statistic for the gene-set *H* is the maximum association score of any non-empty subset of *H* i.e. 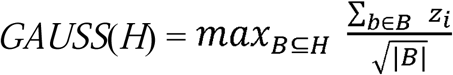, where |*B*| is the number of genes in the subset. Such maximum type statistics have been used in the context of multiple phenotype test, meta-analysis ^26^, and gene-environment interaction tests ^27^. Although we consider the maximum over all 2^*m*^−1 non-empty subsets of *H*, in practice the GAUSS test statistic can be obtained through an algorithm with the computational complexity of *O*(*m*log*m*) (See Methods). We term the subset of genes *B* for which the maximum is attained the **core subset** (CS) of the gene-set *H*.

#### Step 2, p-value calculation

Due to LD between variants in genes in the same genomic region, z-statistics in Step 1 may be dependent. Thus, it is challenging to derive the null distribution of the GAUSS analytically. Instead, we employ a fast simulation approach. We first estimate the correlation structure 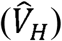 among the z-statistics (*z*_1_, *z*_2_, *…, z*_*m*_) under the null hypothesis, which can be estimated using the sample itself or an ancestry-matched genotype reference panel. Here we use genotype data from the European individuals in publicly available 1000-Genomes data ^28^ as the reference panel. We note that 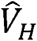 needs to be estimated only once for a given dataset and reused for all iterations. With 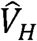, we approximate the joint distribution of z-statistics using a multivariate normal distribution. Now the null distribution of GAUSS test statistics can be simulated by repeatedly generating z-statistics from the mean zero multivariate normal distribution with covariance 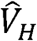 and calculating GAUSS statistics from the simulated sets of z-statistics, with the proportion of simulated null test statistics greater than the observed GAUSS statistic an estimate of the p-value (See Methods). To further reduce computational cost, we use an adaptive resampling scheme (See Methods). To estimate very small p-values (e.g. p-value < 5×10^−6^), we use a generalized Pareto distribution (GPD)-based method ^18^ (See Supplementary Section A). We fit a GPD to the upper tail of the simulated GAUSS test statistics using right-tailed second order Anderson-Darling statistic (GPD-AD2R) (See Supplementary Section A and Supplementary Figure 1) and estimate the p-value by inverting the distribution function of the fitted GPD.

### Simulation results

We carried out simulation studies to evaluate the performance of GAUSS. To understand the effect of the number of genes in the gene-sets, we selected three gene-sets of varying length from GO terms in MSigDB (v6.2) for our simulations: regulation of blood volume by renin angiotensin (GO: 0002016; 11 genes), sterol metabolic process (GO: 0016125; 123 genes) and immune response process (GO: 0006955; 1100 genes).

#### Type I error rates and power to identify associated gene sets

To reflect realistic LD-patterns, we used genotypes of 5,000 unrelated UK Biobank participants throughout our simulations. To estimate the type-I error, we generated a normally distributed phenotype for these same individuals (See Simulation Model), independent of genotypes. We then calculated the gene-based p-values using SKAT-Common-Rare test for each gene in the gene-sets and subsequently applied the GAUSS test. Type-I errors of GAUSS remained well calibrated at *α* = 1 × 10^−4^, 1 × 10^−5^ and 5 × 10^−6^ (Table 1) for all three gene-sets.

**Table 1:** Estimated type-I error of GAUSS for gene-sets GO: 0016125, GO: 0006955 and GO: 0002016.

| $\alpha$ | GO: 0016125<br>(123 genes) | GO: 0006955<br>(1100 genes) | GO: 0002016<br>(11 genes) |
| --- | --- | --- | --- |
| $1 \times 10^{-04}$ | $9.8 \times 10^{-05}$ | $9.8 \times 10^{-05}$ | $9.7 \times 10^{-05}$ |
| $1 \times 10^{-05}$ | $9.9 \times 10^{-06}$ | $9.3 \times 10^{-06}$ | $9.6 \times 10^{-06}$ |
| $5 \times 10^{-06}$ | $4.6 \times 10^{-06}$ | $4.8 \times 10^{-06}$ | $4.2 \times 10^{-06}$ |

Next, we compared the power to detect a gene set-phenotype association, under a spectrum of association models for GAUSS and three existing methods: SKAT for all the variants in the gene-set (SKAT-Pathway), MAGMA, and aSPUpath. With gene-set GO: 0016125, we first considered a scenario that 20 of the 123 genes (16.2%) to be active and within each active gene we set 30% of the variants to be causal. We varied the gene-set heritability (*h*_*gs*_^*2*^) from 1% to 6%. The empirical power of each method increased with increasing *h*_*gs*_ ^*2*^. GAUSS and MAGMA had similar power (Figure 1; Left panel) for all scenarios, and SKAT-Pathway had the lowest power. aSPUpath had slightly lower power than GAUSS when *h*_*gs*_ ^*2*^ *=* 1-3% and had similar power for *h*_*gs*_ ^*2 =*^ 4-6%.

**Figure 1.**
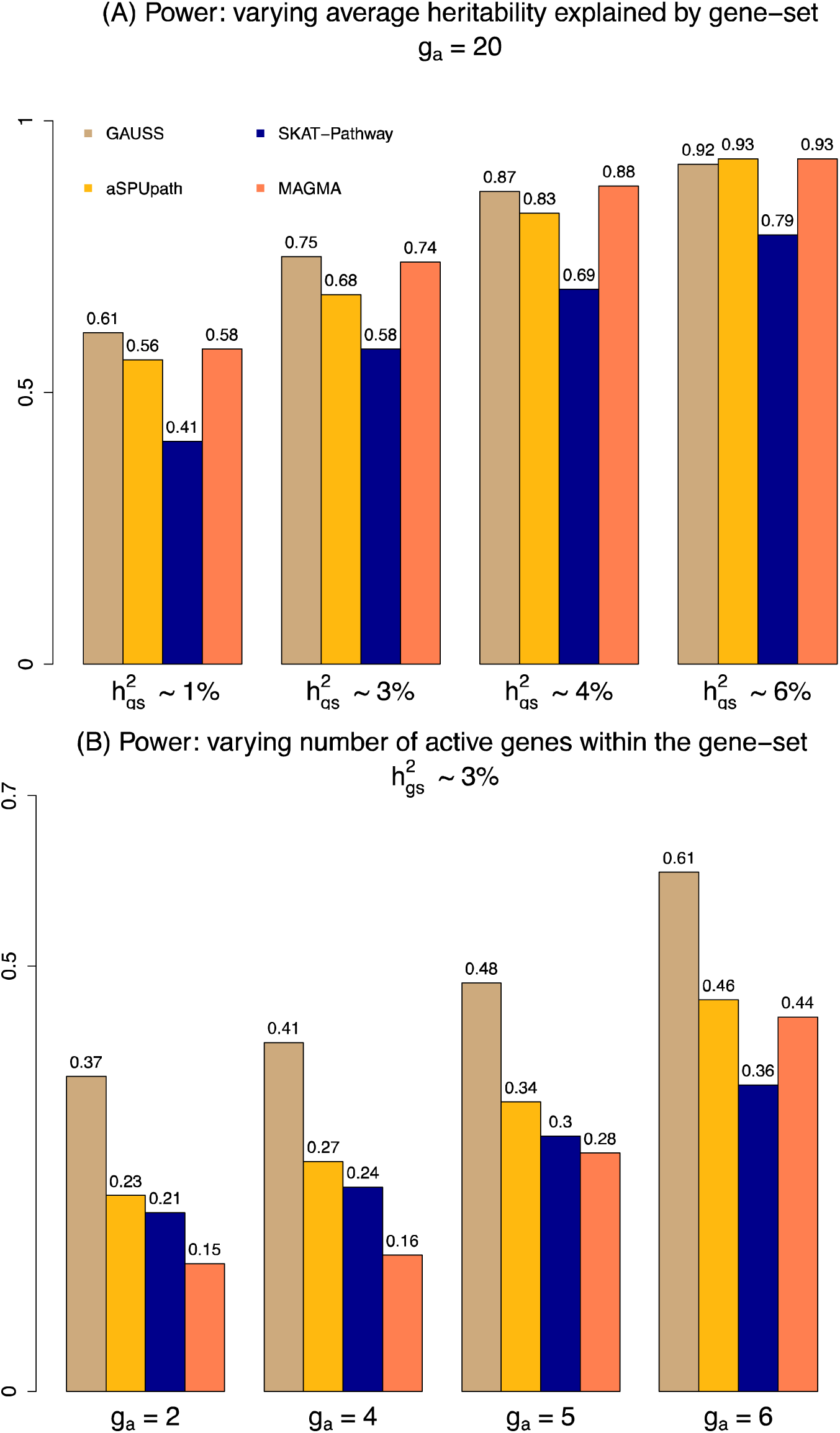
Empirical power for GAUSS. Estimated power of GAUSS using GO: 0016125 gene-set (123 genes), compared with that of aSPUpath, SKAT-Pathway and MAGMA under different average heritability explained (*h*_*gs*_^*2*^) and different number of active genes (*g*_*a*_). (a) Power of GAUSS when 20 genes are active (*g*_*a*_ = 20) and the variants with different average heritability (*h*_*gs*_^*2*^) explained by the gene-set. (b) Power of GAUSS with different number of active genes (2, 4, 5 and 6) and the gene-set has an average heritability *h*_*gs*_^*2*^ of 3%. The proportion of causal variants in an active gene (See Simulation Model) was set to be 30%.

Next, we considered a scenario where the signals were sparser (Figure 1; Right panel), i.e., 2 (1.6%) to 6 (5.0%) genes among the 123 genes in the gene-set were active. We fixed the gene-set heritability (*h*_*gs*_ ^*2*^) at ∼3%. In all the simulation settings, GAUSS was the most powerful method. The power gap between GAUSS and the other methods was particularly large when only 2 genes were active. Among the other methods, aSPUpath had the second highest power and MAGMA had the lowest power when 10 to 20 genes were active. The overall trend remained similar when we used a much larger gene-set like GO: 0006955 or a much smaller gene-set like GO: 0002016 (Supplementary Figures 2 and 3).

#### Identification of active genes

We investigated the sensitivity and specificity of GAUSS in identifying active genes through the core subset (CS) genes. Sensitivity is defined as the proportion of active genes correctly identified by GAUSS as CS genes, specificity as the proportion of inactive genes correctly identified by GAUSS as not CS genes. Since no current methods attempt to identify the active genes within the gene-set, we compared the performance of GAUSS to the heuristic approach of defining the significant genes (p-value < 2.5×10^−6^) as active. For GAUSS, both sensitivity and specificity remained higher (>75%) than the significant genes approach at different values of *h*_*gs*_^*2*^ and for varying number of active genes (Figure 2). We also evaluated power to identify the exact set of active genes which is a more stringent criteria compared to sensitivity and specificity. Under different magnitudes of gene effect sizes defined by different values of heritability, the empirical probability to identify the exact set of active genes via GAUSS had a slight decreasing trend with increasing number of active genes (Figure 2). The overall patterns remained similar when we varied the length of the gene-set by using GO: 0006955 (1100 genes) and GO: 0002016 (11 genes) for our simulations (See Supplementary Figure 4-5).

**Figure 2.**
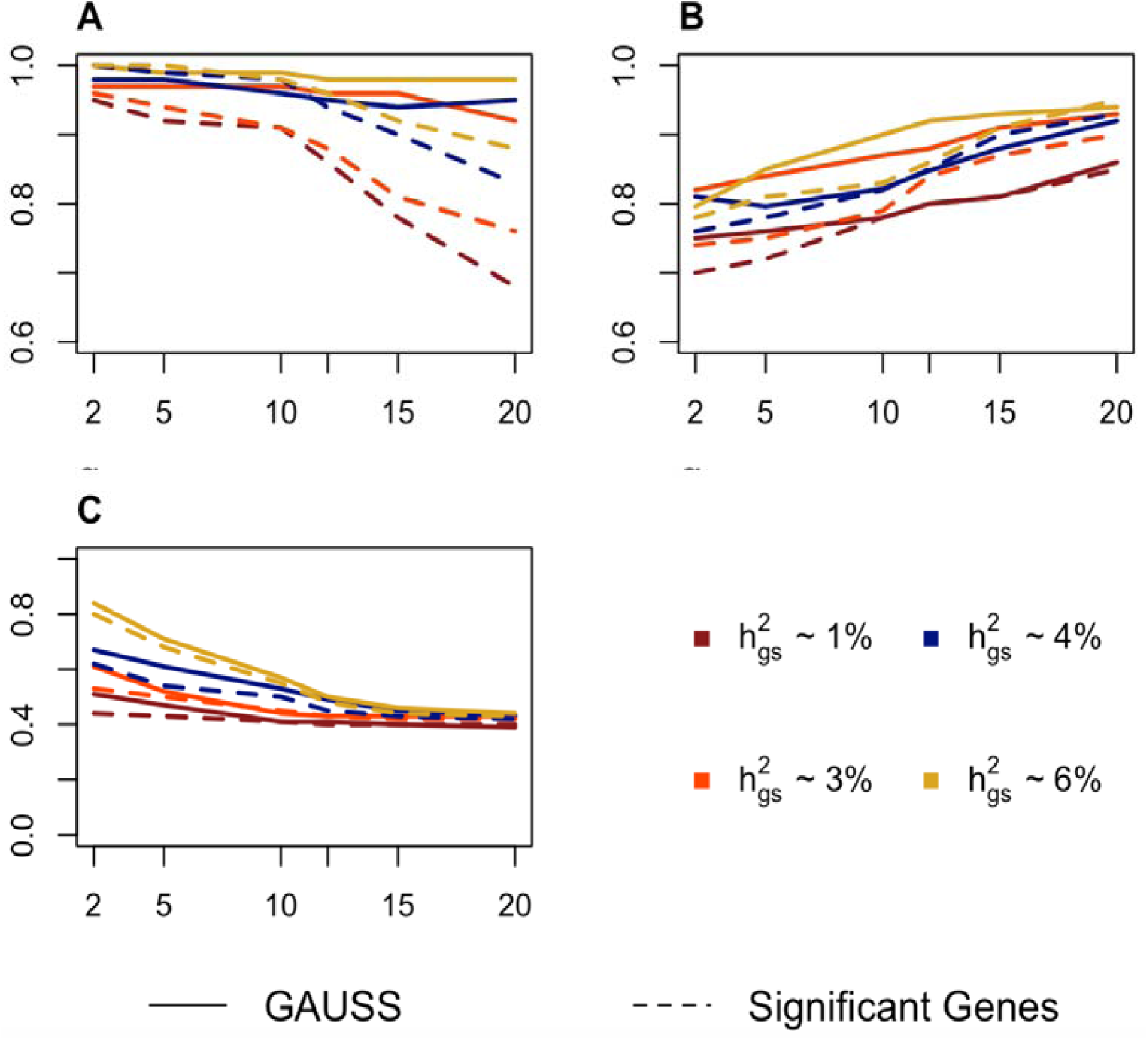
Sensitivity, Specificity and probability of identifying the exact non-null subset for GAUSS. The results are across different number of active genes in the gene-set (horizontal axis) and different average heritability explained by the gene-set (*h*_*gs*_^*2*^) for GO: 0016125 (123 genes). (A) Sensitivity, (B) Specificity and (C) Probability of identifying an exact non-null subset using GAUSS (solid line) compared with the method of using the set of significant genes as the active gene set (dashed line). The proportion of causal variants in an active gene (See Simulation Model) was set to be 30%.

Simulation results highlight the utility of GAUSS compared to the existing methods. Especially when only a few genes in the gene-set are weakly associated with the phenotype, GAUSS has greater power to identify gene-set associations. Further by extracting CS genes, GAUSS can identify the set of active genes with high probability and provides a direct way to interpret findings.

### Association analysis in UK Biobank

We applied GAUSS to the UK Biobank GWAS summary results for 1,403 binary phenotypes^29^ to identify disease related gene-sets and the corresponding core genes (See Methods). We used 10,679 gene-sets from two MsigDB (v6.2) collections: 1) the curated gene-sets (C2) from KEGG, BioCarta, and Reactome databases and gene-sets representing expression signatures of genetic and chemical perturbations, and 2) gene sets that contain genes annotated by GO term (C5). For each phenotype, we estimated the gene-based (SKAT-Common-Rare) p-value for 18,334 genes using SAIGE summary statistics and LD information from a reference panel consisting of unrelated Europeans in the 1000-Genomes Project (See Methods). For each pair of phenotype and gene-set we computed the GAUSS test-statistic, corresponding p-value, and the core subset (CS) of genes (if the gene-set is reported to be significant). We used the Bonferroni corrected gene set p-value threshold for each phenotype: 0.05/10,679 ≈ 5 × 10^−6^.

#### Overview of UK-Biobank results

The 10,679 gene-sets had median size of 36 genes per gene-set (average: 93.2). 94.2% (17,284 of 18,334) of genes belonged to at least one gene-set. In our analysis, we identified 13,466 significant phenotype-gene-set associations at a p-value cut-off of 5 × 10^−6^. Note that the expected number of p-values < 5 × 10^−6^ under no association across all the phenotypes is approximately 75, so the false discovery rate is 0.004. Among the 1,403 phenotypes, 199 (14.1%) had at least one significantly associated gene-set while among the 10,679 gene-sets, 34.1% (3,638) had at least one significantly associated phenotype. There was no significant enrichment in the proportion of association by category of gene-sets, i.e. the GO (C5) gene-sets or Curated (C2) gene-sets (p-value = 0.13). For the significant associations, the average number of the extracted CS genes was 17.2, and a large proportion of the associations (53.6%; 7,237) were due to effects of a single gene within the gene-set. However, 24.6% of the associations were driven by a set of 5 or more CS genes. Approximately 32.7% of the significant associations were with gene-sets that do not have any genes significant at the gene-based cutoff of 2.5× 10^−6^. This underlines that GAUSS can effectively aggregate weaker associations detect significant gene-sets associated to phenotype. Among the different categories of phenotypes, “endocrine/metabolic” diseases had the highest number of associations (5,015; 37.2%), followed by “circulatory system” diseases (2,312; 17.2%) and “digestive” diseases (1,985; 14.7%). (See Supplementary Figures 6-9)

#### Gene-set association analysis for two exemplary phenotypes

To demonstrate the utility of GAUSS in detecting weak associations and improving interpretation, we show association results for two example phenotypes: E.Coli infection (EC; PheCode: 041.4) and Gastritis and duodenitis (GD; PheCode: 535). Single variant GWAS results using SAIGE for these phenotypes can be visualized on UK-Biobank PheWeb (URL: http://pheweb.sph.umich.edu/SAIGE-UKB/pheno/041.4) and do not show any evidence of substantial inflation (λ_*GC*_ varies from 0.91 to 1.09). In the single variant analysis, EC has no genome-wide significant locus and GD has five genome-wide significant loci. When we estimated the gene-based (SKAT-Common-Rare) p-values for EC and GD, the QQ plots were well calibrated without any indication of inflation (λ_*GC*_ varies from 0.98 to 1.01; Supplementary Figure 10). At a gene-based cut-off of 2.5 × 10^−6^, EC does not have any significantly associated genes; GD has three genes that are significantly associated: *HLA-DQA1* (p-value = 9.8 × 10^−11^), *HLA-DQB1* (p-value = 1.4 × 10^−8^) and *PBX2* (p-value = 2.1 ×10^−6^).

Next, we performed gene-set association analysis using GAUSS (Figure 3). We found that EC is associated with two gene-sets (Figure 3; Left panel): fatty acid catabolic process (GO: 0009062; p-value < 1×10^−6^) and fatty acid beta oxidation (GO: 0006635; p-value = 2 × 10^−6^). Although a thorough gene-set association analysis of E.Coli infection has not been done before to our knowledge, the antibacterial role of fatty acids has been well-reported ^30–32^. A set of 25 distinct genes (Table 2) is selected by GAUSS as the CS genes that are responsible for the association although none of them are marginally associated with EC (minimum p-value= 2.2 × 10^−4^), demonstrating that GAUSS can effectively aggregate weaker signals within a gene-set, which would otherwise not have been detected.

**Table 2:**
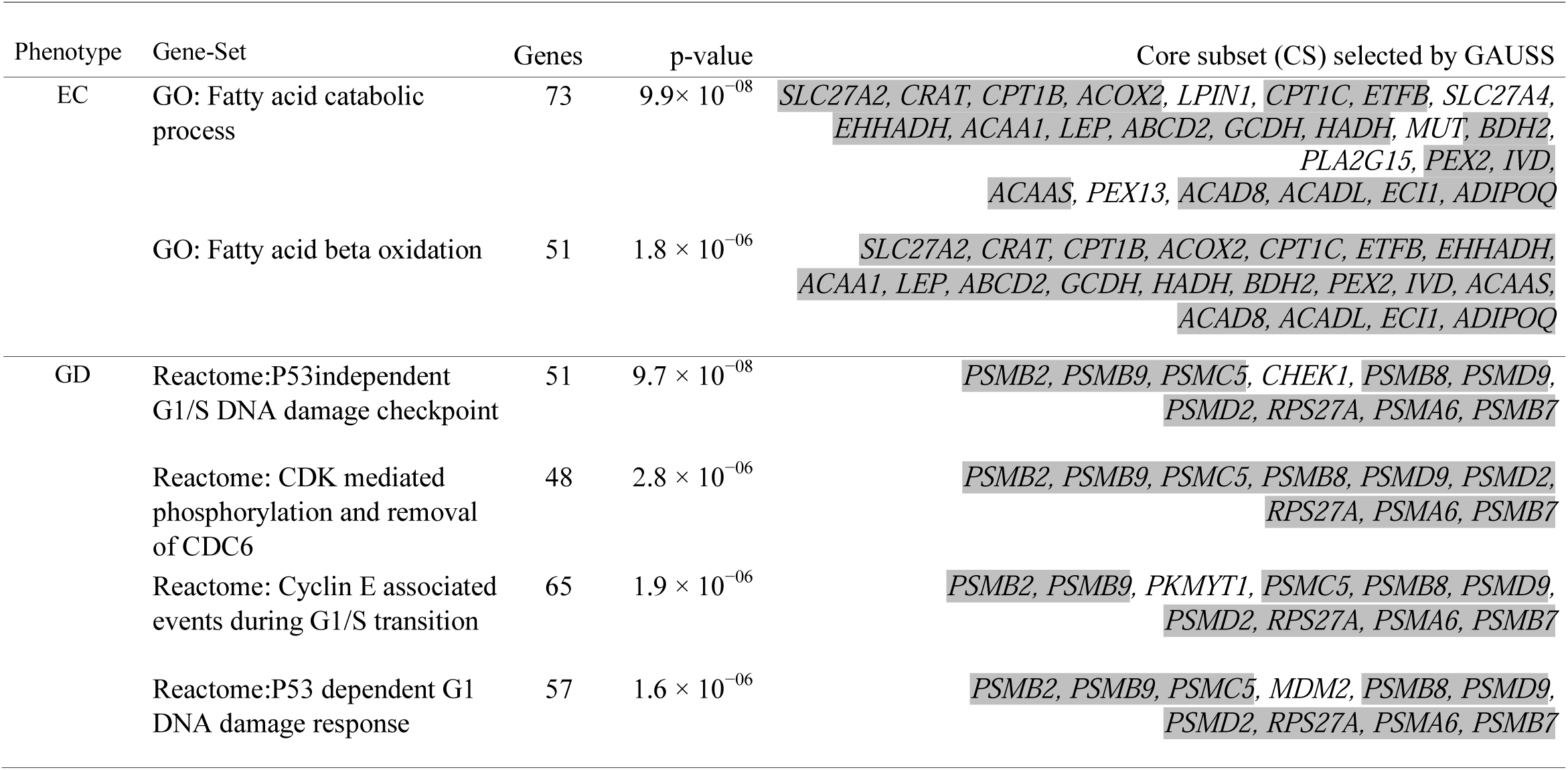
Significant gene-sets associated with E. Coli infection (EC) and Gastritis and duodenitis (GD) corresponding p-values and the CS genes selected by GAUSS. Gray-shade genes are shared CS genes in different pathways for a phenotype. P-values < 1 × 10^−06^ were estimated using GPD (See Results: Overview of methods).

**Figure 3.**
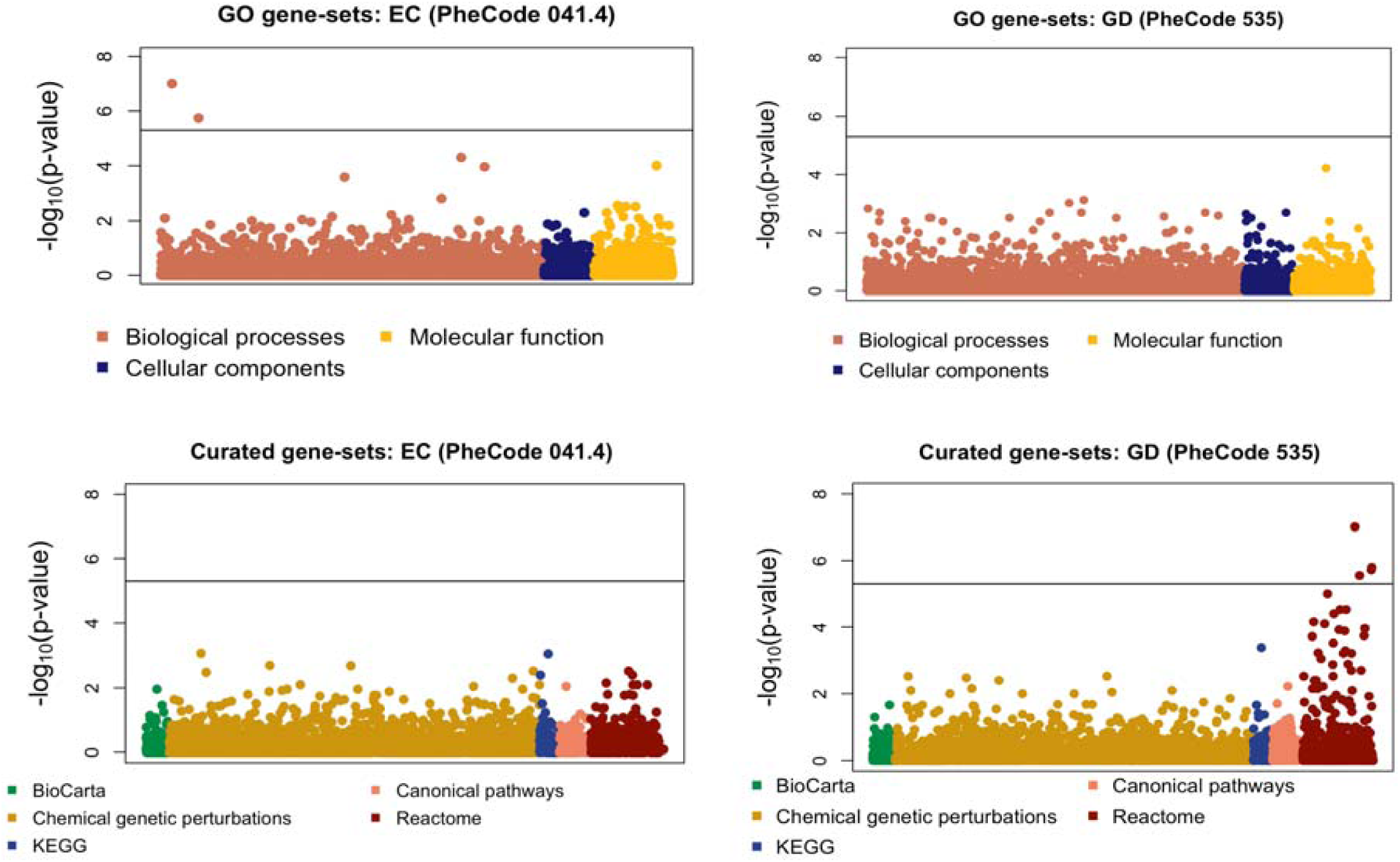
P-values for EC and GD across GO and Curated gene-sets. P-values for association of E. Coli infection (EC; PheCode 041.4) and Gastritis and duodenitis (GD; PheCode 535) with the GO pathways (C5; Upper Panel) and Curated pathways (C2; Lower Panel). P-values < 1 × 10^−06^ were estimated using GPD (See Results: Overview of methods). The horizontal solid black line denotes the significance threshold of 5 × 10^−06^.

In gene-set association analysis of GD (Figure 3; Right panel), we found 4 gene-sets to be associated (Table 2). Although the gene-sets and the corresponding functions are biologically related, their role in GD is not easily identifiable. GAUSS selects a set of 10 genes to be the CS genes for the gene-sets, the majority being from the different proteasome endopeptidase complex (*PSM*) subunits. Different proteasome subunit genes have been found to be associated with several inflammatory responses and intestinal diseases^33,34^. In particular, the role of *PSMB8* ^35^ in gastric cancer has been extensively reported in the literature. Also, *PSMB9* and *PSMB8* have been found to be associated with several gastrointestinal disorders like celiac disease and inflammatory bowel disease ^36–38^. Although, none of these genes are individually significantly associated with GD (minimum p-value=2.8 × 10^−4^), they jointly drive the strong association signal. This highlights that the selected core genes (CS) can help in finding biological targets for downstream investigation.

To validate the results, we tested the gene-sets identified by GAUSS for EC and GD (Table 2) in an independent dataset. For this, we applied GAUSS to summary data from the Michigan Genomics Initiative (MGI) of about 38,000 European samples^39^ (See Supplementary Section B). Our results show that 5 out of the 6 gene-sets that were significant in UK-Biobank for either of EC or GD, had a nominal evidence of significance (p-value < 0.05; Supplementary Table 1). Given that the sample size of MGI is about 10 times lower than UK Biobank, our findings indicate that the detected associations are potentially true.

#### Phenome-wide association analysis for single gene-set

We further analyzed the association of a gene-set across the binary phenome. Figure 4 shows association results across the 1,403 phenotypes for one example gene-set: ATP-binding cassette (ABC) transporters from KEGG (ABC transporters; URL). ABC transporters are involved in tumor resistance, cystic fibrosis, and a spectrum of other heritable phenotypes along with the development of resistance to several drugs^40^. We found 18 phenotypes significantly associated (p-value < 5×10^−6^) with ABC transporters (Table 3), mainly from “digestive” disease and “endocrine/metabolic” disease categories. Among the CS genes selected for different associated phenotypes, *TAP2* is the most frequent. *TAP2* has been reported to be associated with several phenotypes including diastolic blood pressure ^41^, type-1 diabetes, and autoimmune thyroid diseases ^42^. Our results suggest that the significant association of ABC transporters to disorders like psoriasis, celiac disease, and type-1 diabetes are mainly driven by the single-gene effect of *TAP2*. However, the association of ABC transporters with gout, lipoid metabolism, and gallstones are driven mainly by *ABCG5* and *ABCG2*. Thus, although ABC transporters gene-set is significantly associated with 18 phenotypes, the CS genes that drive the associations are different, which can be indicative of different mechanisms underlying the phenotypes.

**Table 3:** Phenotypes associated with ABC transporters gene-set, corresponding p-values and the CS genes selected by GAUSS. P-values < 1 × 10^−06^ were estimated using GPD (See Results: Overview of methods).

| Phenotype | Category | PheCode | p-value | Core subset (CS) selected by GAUSS |
| --- | --- | --- | --- | --- |
| Psoriasis | Dermatologic | 696.4 | $9.1 \times 10^{-11}$ | *TAP2* |
| Psoriasis and related disorders | Dermatologic | 696 | $3.5 \times 10^{-11}$ | *TAP2* |
| Celiac disease | Digestive | 557.1 | $1.8 \times 10^{-32}$ | *TAP2* |
| Intestinal malabsorptions (non-celiac) | Digestive | 557 | $3.4 \times 10^{-31}$ | *TAP2* |
| Cholelithiasis with other cholecystitis | Digestive | 574.12 | $1.8 \times 10^{-19}$ | *ABCG5* |
| Cholelithiasis | Digestive | 574.1 | $1.2 \times 10^{-31}$ | *ABCG5* |
| Calculus of bile duct | Digestive | 574.2 | $1.2 \times 10^{-11}$ | *ABCG5* |
| Cholelithiasis without cholecystitis | Digestive | 574.3 | $4.1 \times 10^{-06}$ | *ABCG5, ABCC12, ABCA8, ABCB4* |
| Cholelithiasis and cholecystitis | Digestive | 574 | $9.6 \times 10^{-34}$ | *ABCG5* |
| Other biliary tract disease | Digestive | 575 | $6.4 \times 10^{-08}$ | *ABCG5* |
| Hypothyroidism NOS | endocrine/metabolic | 244.4 | $1.5 \times 10^{-12}$ | *TAP2* |
| Type-1 diabetes | endocrine/metabolic | 250.1 | $2.9 \times 10^{-08}$ | *TAP2* |
| Hypercholesterolemia | endocrine/metabolic | 272.11 | $2.8 \times 10^{-08}$ | *ABCG5, TAP2, ABCC10, ABCA2, ABCA5, ABCA1, ABCA6, ABCC12, ABCC1, ABCA8, ABCB9* |
| Hyperlipidemia | endocrine/metabolic | 272.1 | $2.1 \times 10^{-07}$ | *TAP2, ABCG5, ABCC10, ABCA6, ABCA2, ABCA5, ABCA1, ABCC1, ABCA8* |
| Disorders of lipid metabolism | endocrine/metabolic | 272 | $5.7 \times 10^{-27}$ | *ABCG2* |
| Gout | endocrine/metabolic | 274.1 | $7.1 \times 10^{-10}$ | *ABCG2* |
| Gout and other crystal arthropathies | endocrine/metabolic | 274 | $3.8 \times 10^{-06}$ | *ABCG2* |
| Asthma | Respiratory | 495 | $9.1 \times 10^{-11}$ | *TAP2* |

**Figure 4.**
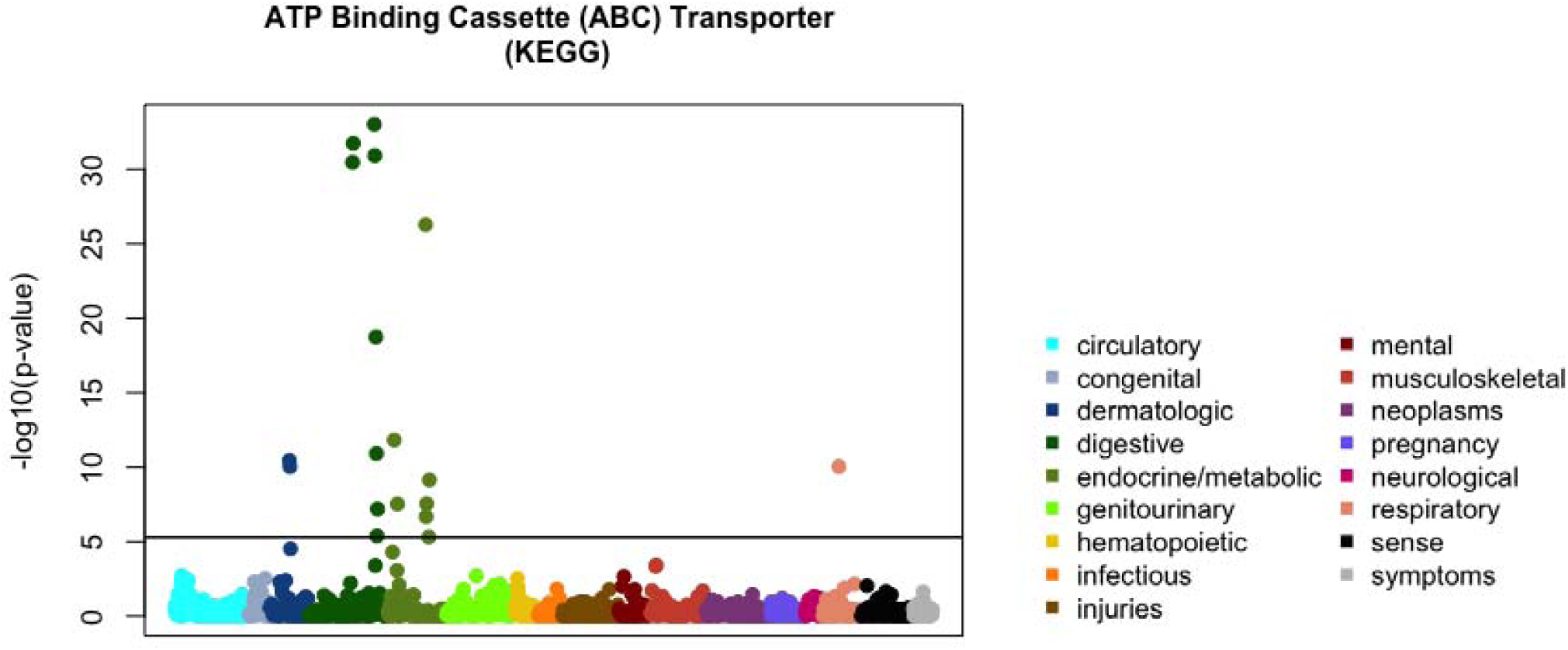
Phenome-wide p-values for ABC Transporter pathway in KEGG: P-values for association of 1403 phenotypes with ABC transporters pathway (KEGG). The horizontal solid black line denotes the significance threshold of 5 × 10^−06^. P-values < 1 × 10^−06^ were estimated through the GPD method (See Results: Overview of methods).

### Computation time comparison

GAUSS uses an adaptive resampling scheme to estimate the p-value (See Methods). Hence, the computation time of GAUSS can vary across different phenotypes depending on the number of associated gene-sets. To evaluate the computation time of GAUSS in phenotypes with small and large number of associated gene-sets, we chose two phenotypes in UK Biobank data: Pernicious anemia (PA; PheCode: 281.11), which had only 1 associated gene-set and type-2 diabetes (T2D; PheCode: 250.2) which had 227 associated gene-sets. Supplementary Figure 11 shows the total run-time (in CPU-hours) of GAUSS, MAGMA, and aSPUpath for UK Biobank. Total run-times were calculated as the net time taken starting from the input of summary statistics until the p-values for 10,679 gene-sets were generated. In terms of total run-time, MAGMA (8.1 and 8.3 CPU-hours for PA and T2D respectively) performed slightly better than GAUSS (10.3 and 12.8 CPU-hours respectively). aSPUpath (93 and 98 CPU-hours respectively) was substantially slower than all other methods. For the full UK Biobank analysis, GAUSS had an average run-time of 11.2 CPU-hours per phenotype.

To obtain GAUSS p-values, we need to estimate the null correlation structure 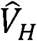. 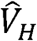 needs to be calculated once for a given dataset and can be used for the analysis across all different phenotypes. We calculated 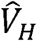 using a reference panel consisting of the unrelated individuals of European ancestry in 1000-Genomes dataset. This required 723 CPU-hours. However, since the calculation can be parallelizable, the actual clock-time was around 12 hours with 60 CPU-cores.

## Discussion

Here we present GAUSS, which uses a subset-based statistic to test the association between a gene-set and a phenotype. Similar to several existing approaches like MAGMA and aSPUpath, GAUSS aims to aggregate weak to moderate association signals across a set of genes. Additionally, GAUSS identifies the core subset (CS) of genes which maximize the association signals.

The identification of CS genes within a gene-set is a key feature of GAUSS. To the best of our knowledge, there exists no gene-set method to adaptively identify the subset of genes that drives the association signal. Most existing approaches suggest using the genes with the lowest p-values in the gene-set. In contrast, GAUSS selects CS genes that have the maximum association score. The selected CS genes can highlight possible underlying mechanisms and can be used for downstream analysis. Furthermore, the association results for a given gene-set across many phenotypes can highlight the underlying biological similarities or differences between phenotypes, especially through the CS genes.

GAUSS uses several approximations and adaptive approaches to reduce computation cost. It summarizes the gene-based association signals to z-scores and uses Gaussian copula to model the joint distribution among z-scores. This approach allows estimation of the distribution of GAUSS statistics by generating multivariate normal random variables and is far more efficient than using standard permutation approaches. In addition, GAUSS uses adaptive resampling methods and GDP-based small p-value estimation, which further reduces computation cost.

In analysis of simulated and UK-Biobank data, we used European ancestry samples of 1000-Genomes data for reference data. To investigate whether the method is sensitive to the reference panel, we compared the performance of GAUSS using 1000-Genomes data to that using UK-Biobank data as reference (Supplementary Figure 12). The results show that the choice of reference panel did not substantially impact the results from GAUSS. However, it is important that the reference population and the study sample belong to the same ancestry to reflect similar LD patterns.

Our UK Biobank analysis shows that typically only a small percentage of genes in the pathway are selected as core genes (Supplementary Figure 8). Simulations show that GAUSS has substantially greater power than the existing methods in detecting associations in such sparse scenarios. Existing methods, such as MAGMA, use test statistics that are averaged over all the variants or genes in the gene-set. If the fraction of associated variants is relatively low and associated variants have weak effects, these tests might have low power. However, GAUSS uses a subset-based approach to choose the subset with maximum evidence of association, and thus does not average over all associated and unassociated variants or genes in the gene-set. Hence, even when the fraction of associated variants is low, GAUSS can have greater power. When many of the genes in the gene-set are associated, the power of GAUSS was similar to MAGMA. Thus, in most of the practical scenarios GAUSS has power greater than or equal to that of existing methods MAGMA and aSPUpath. Further, the type-I error for GAUSS remains calibrated at the desired level.

A limitation of GAUSS is that it only allows testing for the self-contained null hypothesis. Although this allows us to detect association of a gene-set with a phenotype but does not provide information on enrichment of the associated gene-set. Further, the GPD method of estimating very small p-values needs additional research and exploration.

In summary, we have shown that GAUSS can be more powerful than the existing methods to detect gene-sets associated with phenotypes and facilitates interpretation of gene-set analysis results through CS genes. The novel insights generated by GAUSS and its computational scalability make it an attractive choice to perform phenome-wide gene-set analysis. Our UK Biobank analysis identified large-numbers of gene-set by phenotype association pairs, and we have partially validated associations in EC and GD phenotypes through MGI data analysis. By providing powerful, scalable, and more interpretable gene-set analysis results, our approach will contribute to identifying genetic components of complex phenotypes. We have made a GAUSS software package and UK-Biobank analysis results publicly available (See URL).

## Acknowledgements

This research was supported by grants R01-HG008773 and R01-LM012535 (D.D. and S.L.) and R01-HG009976 (M.B.). UK Biobank data was accessed under the accession number: 45227. The authors acknowledge the Michigan Genomics Initiative participants, Precision Health at the University of Michigan, the University of Michigan Medical School Central Biorepository, and the University of Michigan Advanced Genomics Core for providing data and specimen storage, management, processing, and distribution services, and the Center for Statistical Genetics in the Department of Biostatistics at the School of Public Health for genotype data curation, imputation, and management in support of the research reported in this publication.

## URL

GAUSS: https://github.com/diptavo/GAUSS

PathWeb: http://ukb-pathway.leelabsg.org/

MsigDB (v6.2): https://data.broadinstitute.org/gsea-msigdb/msigdb/release/6.2/

UK-Biobank single variant analysis Pheweb: http://pheweb.sph.umich.edu/SAIGE-UKB/

emeraLD: https://github.com/statgen/emeraLD

## Methods

### Estimating gene-based p-values from summary statistics

Let *y* = (*y*_1_,*y*_2_, …,*y*_*n*_)^*T*^ be the vector of phenotype for n individuals; *X* the matrix of q non-genetic covariates including the intercept; *G*_*j*_ = (*G*_1*j*_,*G*_2*j*_, …, *G*_*nj*_)^*T*^ the vector of the minor allele counts (0, 1, or 2) for genetic variant j; and *G* = (*G*_1_,*G*_2_, …, *G*_*m*_)^*T*^ the genotype matrix for *m* genetic variants in a target gene or region. The regression model used to relate the phenotype to the *m* genetic variants in the region is:

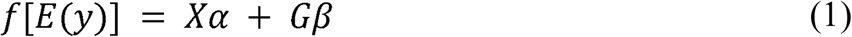

where *f*(.) is a link function and can be set to be the identity function for continuous traits or the logistic function for binary traits, *α* is the vector of regression coefficients of *q* non-genetic covariates; and *β* = (*β*_1_,…, *β*_*m*_) ^*T*^ is the vector of regression coefficients of the *m* genetic variants. To test *H*_0_: *β* = 0, under the random effects assumption *β*_*i*_ *∼ N* (0, τ ^2^), the SKAT test statistic [35] is

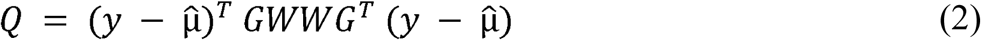

where 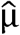 is the estimated expected value of *y* under the null hypothesis of no association and *W* = *diag*(*w*_1_,.., *w*_*m*_) is a diagonal weighting matrix. Wu et al ^22^ suggested to use the *Beta (MAF, 1,25)* density function as a weight to upweight rarer variants. Under the null hypothesis, *Q* asymptotically follows a mixture of chi-squared distributions and p-values can be computed by inverting the characteristic function. The mixing parameters are the eigenvalues of *WG*^*T*^ *P*_0_*GW*, where *P*_0_ = *I*_*n*_ *− X*(*X*^*T*^ *X*)^−1^*X*^*T*^ and *I*_*n*_ is the identity matrix of order *n*.

Equation (1) uses individual level data on the samples. However, the test of association can be effectively approximated by using summary statistics on the *m* variants in the region ^25,43^. Given the estimated GWAS summary statistics (*MAF*_*j*_, *β*_*j*_, *SE*_*j*_) for each variant j, the test statistic *Q* in Equation (2) can be approximated as

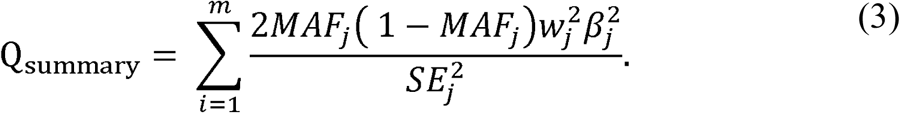

Under the null hypothesis, *Q* follows a mixture of chi-squares and the mixing parameters are the eigenvalues of the matrix *WG*^*T*^ *P*_0_*GW*. Replacing *P*_0_ by Φ_0_ = *I −* 11^*T*^ */n*, we can approximate the eigenvalues by that of the matrix *WG*^*T*^ Φ_0_*GW*. The matrix *G*^*T*^ Φ_0_*G* is the LD-matrix of the *m* variants, which can be estimated using a publicly available reference panel ^28^.

Ionita-Laza et al. ^23^ developed SKAT-Common-Rare which tests the combined effect of rare and common variants in a region. Given summary statistics as above, we construct the test statistic separately for common and rare variants as *Q*_*summary*; *common*_ and *Q*_*summary; rare*_ using equation (3). For common variants, we used the *Beta*(MAF, 0.5,0.5) density function for weight calculation, and for rare variants, Beta(MAF, 1, 25) ^23^. Then, SKAT-Common-Rare test is then constructed as

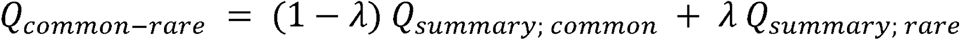

where 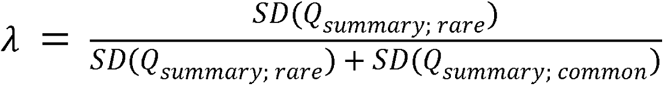. The asymptotic null distribution of Q_common-rare_ is a mixture of chi-squares and can be approximated using the empirical LD matrices of common and rare variants.

### GAUSS test statistic

Suppose *Pvalue*_*i*_ is the p-value of the *i*^*th*^ gene (i=1, …, m) in the gene-set *H*. We first convert the *Pvalue*_*i*_ to a z-score as *z*_*i*_ = Φ^−1^ (1 − *Pvalue*_*i*_), where Φ^−1^ is the inverse of the standard normal cumulative distribution function. Here we have used SKAT-Common-Rare to obtain *Pvalue*_*i*_ from GWAS summary statistics, but other tests like prediXcan can be used to obtain gene-based p-values.

For any non-empty subset *B* ⊆ *H*, we define S(B) the association score for the subset B as 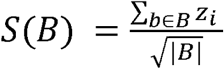 where |*B*| is the number of genes in *B*. We define the GAUSS statistic for the gene-set *H* as the maximum score of a subset of *H*

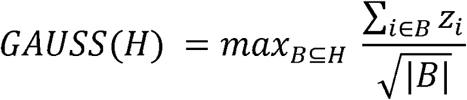

Although the maximum is over all 2^*m*^ − 1 possible non-empty subsets of *H*, the computational complexity can be greatly reduced by rewriting the formula as,

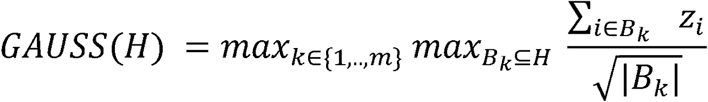

where *B*_*k*_ denotes a non-empty subset of *H* with *k* elements. It is easy to show that

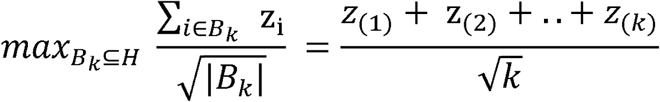

where z_(1)_, z_(2)_, …, z_(m)_ are the ordered z-statistic in decreasing order with z_(1)_ being the maximum. We implement the following algorithm to obtain the GAUSS statistic as:

1. Order the z-statistic for the m genes as z_(1)_, z_(2)_, …, z_(m)_
2. Starting with *k* =1 compute 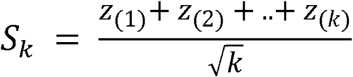 for all k = 1, 2, …, m
3. Calculate the GAUSS test statistic as *max*_*k* ∈ {1, *…,m*}_ *S*_*k*_

Using this approach, computational cost is reduced from *O(2*^*m*^) to *O(mlogm)*.

### Fast estimation of the p-value of GAUSS

We employ a fast two-step approach which uses a normal-Copula to estimate p-values for GAUSS. We first estimate the correlation structure 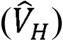 among the z-statistics z_1_, z_2_, …, z_m_ under the null hypothesis of no association through a small number of simulations using a reference LD panel (see next section). Then we estimate the p-value of the GAUSS test statistic as follows:

1. Starting from *r* = 1, in the *r*^*th*^ step, generate a random *m* vector *Z*_*r*_ from the multivariate normal distribution 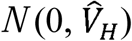
2. Calculate the GAUSS statistic using *Z*_*r*_ as above, *GAUSS*(*H*)_*r*_
3. Repeat steps 1 and 2 R times, say R (= 10^6^)
4. Estimate the p-value for the observed *GAUSS*(*H*)_*r*_ as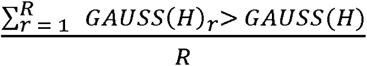

Although it is a simulation-based method, the algorithm can be efficiently implemented since it only requires generating multivariate normal (MVN) random vectors. For example, generating 1 million MVN random vectors for a gene-set with 100 genes (*m* = 100) requires 2 CPU-seconds on an Intel Xeon 2.80 □ GHz computer.

We also implemented an adaptive resampling scheme that performs fewer iterations if the p-value is large (say >0.005). For a given GAUSS test statistic, we first use 1000 iterations to estimate the p-value. If the estimated p-value is ≤ 0.005, then we perform 10^6^ iterations to more accurately estimate the p-value. Thus, if the true p-value is large (> 0.005) the above algorithm estimates it in less than 1 CPU-second and if the true p-value is small the algorithm takes 161 CPU seconds on average. If the true p-value is very small, 10^6^ resampling cannot estimate it. For this, we use GPD based approximation approach (Supplementary Section A).

### Reference data and the estimation of correlation structure *V*_H_

Given the GWAS summary statistic for a phenotype, to obtain the GAUSS p-value for a gene-set, we have used the reference panel twice. First, we used the reference panel to extract LD across variants in a gene or region. This LD information is used to construct the null distribution and evaluate the gene-based p-value. We use emeraLD ^44^ (See URL) for fast extraction of LD from variant-call-format (VCF) files. Second, we used the reference panel to estimate the null correlation matrix *V*_*H*_ among the z-statistics. This is a pre-computed matrix which needs to be computed once from the reference data and can be reused for future applications. To estimate this matrix, we generated a null continuous phenotype from the standard normal distribution, computed the gene-based p-values for the annotated genes using SKAT-Common-Rare, and converted them to z-statistics. We repeated this procedure 1000 times and calculated *V*_H_ as the Pearson’s correlation between 1000 null z-statistic values. This approach greatly reduces the computational burden of GAUSS since it does not need to estimate *V*_H_ for every iteration or gene-set.

### Simulation Studies

We used UK-Biobank genotype data for simulation studies. We define a gene within a gene-set as “active” if at least one variant annotated to the gene has non-zero effect size. For a given gene-set we randomly set *g*_*a*_ genes to be active and within the *l*^*th*^ active gene with *t*_*l*_ variants we set *v*_*a*;*l*_ to be the proportion of variants with non-zero effects. Using genotypes of *N* randomly selected unrelated individuals from the UK Biobank we generate the phenotypes for individual *i* (*i* = 1, …, *N*) according to the model

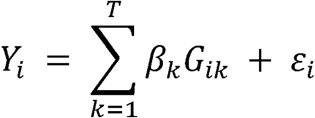

where ε_*i*_ *∼ N* (0,1) and *G*_*ik*_ is the genotype of the i^th^ individual at the k^th^ variant and 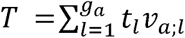 is the total number of variants with non-zero effects. Throughout our simulations, we used *N*=5000. The effect size of the *k*^*th*^ active variant with minor allele frequency *MAF*_*k*_ is generated as *β*_*ik*_ = *c*|*log*_10_(*MAF*_*k*_)| where *c* is the magnitude of the association between a variant and phenotypes. For type-I error simulations we set *c* = 0 while for power we set *c >* 0. We determined the value of *c* by fixing the average heritability explained by the gene-set (*h*_*gs*_ ^*2*^). We used several values for the average heritability explained by the gene-set between *h*_*gs*_ ^*2*^ *=* 1% and 10%. With 20-30% variants having non-zero effect sizes, the corresponding values of *c* varied approximately between 0.10 and 0.25.

With the UK-Biobank genotypes and the simulated phenotypes, we first calculated the GWAS summary statistics for each variant and estimated the gene-based (SKAT-Common-Rare) p-values using the LD extracted from the Europeans in the 1000-Genomes data. Subsequently, we applied GAUSS to the gene-based p-values and extracted the p-value for association. We then calculated power as the fraction of GAUSS p-values less than 5 × 10^−6^ which represents the Bonferroni corrected threshold for testing association across 10,000 independent gene-sets.

### UK-Biobank data analysis

In our GAUSS analysis, we used publicly available UK-Biobank summary statistics that were generated by SAIGE ^29^ (See URL for PheWeb). The summary statistic files included results for markers directly genotyped or imputed by the Haplotype Reference Consortium (HRC) which produced approximately 28 million markers with MAC ≥ 20 and an imputation info score ≥ 0.3. We used EPACTs ^45^ (See URL) with RefSeq gene database for the variant annotation. For each gene we included nonsynonymous variants and variants within 1 kb of the first and last variants in each exon to test for the effect of possibly functional and regulatory variants. We extracted LD information and constructed an LD-matrix from the 1000-Genomes European reference panel using emeraLD. For each of the 1,403 phenotypes and 18,334 genes, we constructed SKAT-Common-Rare test statistic using the estimates from SAIGE of effect size (*β*), standard error (SE), and minor allele frequency (MAF). We transformed the gene-based p-values into z-statistics and performed gene-set analysis for each phenotype.

