## Supplementary for "A powerful subset-based gene-set analysis method identifies novel associations and improves interpretation in UK Biobank"

**Supplementary Materials**

**Section A: Estimation of highly significant p-values**

The statistical significance of GAUSS is calculated as the fraction of null resampling values {*gs_r_*; *r* = 1*,*2*,* ···*, R*} that are at least as large as the original observed GAUSS test statistic (*gs_obs_*). In that case, the minimal p-value that can be observed depends on the number of resampling iterations. For example, if we use 1 million resampling iterations (as used in UK Biobank data analysis), the lowest p-value that can be observed is 1×10^−06^. To estimate smaller p-values (*<* 1×10^−06^), a larger number of resampling iterations are needed which can be computationally expensive. To address this, we have used a tail-based method to accurately estimate smaller p-values using the null resampling values *gs_r_*.

Generalized Pareto Distribution (GPD): In our approach, we model the upper (right) tail of the distribution of *gs_r_* using a generalized pareto distribution which is commonly used for modeling extreme values. In particular, we use the following algorithm similar to Knijnenburg et al^1^ for estimating the p-value:

- Starting with *B* = 250, we fit a generalized pareto distribution to the set {*gs_r_*_(1)_*, gs_r_*_(2)_*,*···*gs_r_*_(_*_B_*_)_}, where *gs_r_*_(1)_ is the maximum value of *gs_r_*
- If the goodness-of-fit p-value is larger than 0.05, we proceed to the next step. Else we reduce *B* by 10 and repeat the previous step.
- Calculate the p-value of *gs_obs_* as $\frac{B}{R} (1- ℘({gs}_{obs}))$, where $℘\left( . \right)$ is the cumulative density function of the fitted generalized pareto distribution.

There are several approaches to fit a generalized pareto distribution to the right tail of the distribution of *gs_r_* ^2–7^. To evaluate the bias and accuracy of these methods in estimating smaller p-values, we performed a small-scale simulation study using different methods for fitting generalized pareto distribution including

1. maximum goodness-of-fit using Kolmogorov-Smirnoff statistic (GPD-KS)
2. maximum likelihood (GPD-MLE)
3. method of moments (GPD-MOM)
4. probability weighted moments (PWM)
5. maximum goodness-of-fit using Anderson-Darling right tail statistic (GPD-ADR)
6. maximum goodness-of-fit using Anderson-Darling right tail statistic of 2^nd^ order (GPD-AD2R)

We used the null correlation structure ($\hat{V}_{H}$) of GO: Cobalamin metabolic process which has 21 genes and set *gs_obs_* = 6. To estimate the simulation-based p-value, we used 10^5^ iterations for a reduced computational burden. Further, we used a smaller number of independent iterations (5 × 10^4^) to estimate the p-value using different GPD methods (Supplementary Figure 1). A visual inspection suggests, GPD-AD2R has low bias and high accuracy (approximate SE= 1. 4×10^−05^) among the methods compared. Based on these results, we use GPD-AD2R as a method to estimate p-values smaller than 1×10^−06^.

**Section B: MGI Data Analysis**:

We did a replication study for the gene-sets that were associated with EC and GD (Table 2) using summary statistic from Michigan Genomics Initiative (MGI). MGI is a biorepository effort to create a longitudinal cohort of participants undergoing anesthesia prior to a surgery or diagnostic procedure, creating a patient community with genome-wide data and electronic health information among others ^8^.

In this replication study, we used summary statistics on 38,009 individuals for approximately 23 million markers that were directly genotyped or imputed using HRC. Using LD information estimated from the unrelated Europeans in 1000 Genomes data we estimated the SKAT-Common-Rare p-values for EC and GD. Following that, we used the gene-sets that were reported in Table 2 to be associated to EC and GD and estimated the corresponding gene-set association p-value using GAUSS.

Among the two gene-sets tested with EC one gene-set reached a nominal significance threshold (p-value < 0.05). For the four gene-sets tested with GD, three reached a nominal threshold (Supplementary Table 1).

Considering the substantially lower number of cases in MGI compared to UK Biobank, this replication study was relatively underpowered (EC: 963 cases; GD: 1926 cases). Despite that, most of the gene-sets associated with EC and GD from the UK Biobank analysis show nominal signals in MGI.

**Supplementary Table 1**: Significant gene-sets associated with E. Coli infection (EC) and Gastritis and duodenitis (GD) corresponding p-values and the CS genes selected by GAUSS in MGI data set.

| Phenotype |  | Gene-set | Genes | p-value  (MGI) | CS genes selected by GAUSS |
| --- | --- | --- | --- | --- | --- |
|  |  | GO: Fatty acid catabolic process | 73 | 0.011 | *ACADS, ETFDH, ABCD2, ECI2, CPT1B, ACAD8, ECI1, PEX2, ACADM, ACADL, CRAT* |
| EC |  | GO: Fatty acid beta oxidation | 51 | 0.029 | *ACADS, ETFDH, ABCD2, LPIN2, CYP4F12, ECI2, CPT1B, ACAD8, ECI1, PEX2, ACADM, ACADL, CRAT, PCCA, CEL* |
|  |  | Reactome: P53 independent  G1/S DNA damage checkpoint | 51 | 0.091 | *RPS27A, PSME1, PSMD6, CDC25A, PSMD2, PSMD13, PSMD7, PSMB2* |
|  |  | Reactome: CDK mediated phosphorylation and removal of CDC6 | 48 | 0.025 | *RPS27A, PSME1, PSMD6, CDC6, PSMD2, PSMD13, PSMD7, PSMB2* |
| GD |  | Reactome: Cyclin E associated events during G1/S transition | 65 | 0.009 | *MYC, RPS27A, PSME1, PSMD6, CDK7, CDC25A, PSMD2, PSMD13, CCNE2, CCNA1, PSMD7, PSMB2* |
|  |  | Reactome: P53 dependent G1  DNA damage response | 57 | 0.044 | *RPS27A, PSME1, PSMD6, PSMD2, PSMD13, CCNE2, PSMD7, PSMB2* |

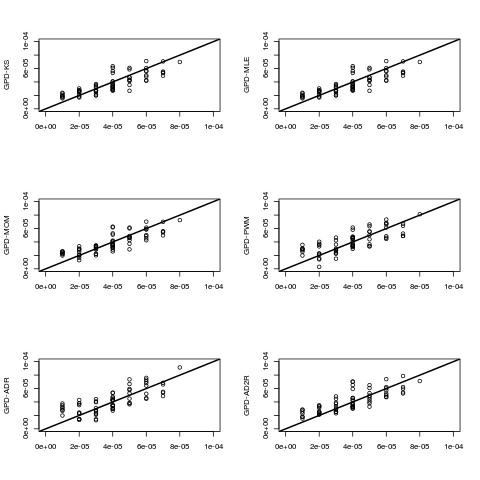

**Supplementary Figure 1**: Bias and accuracy of different GPD methods in estimating small p-values. On the horizontal axis we show the resampling-based p-values plotted against the estimate p-values using different methods of fitting GPD on the vertical axis

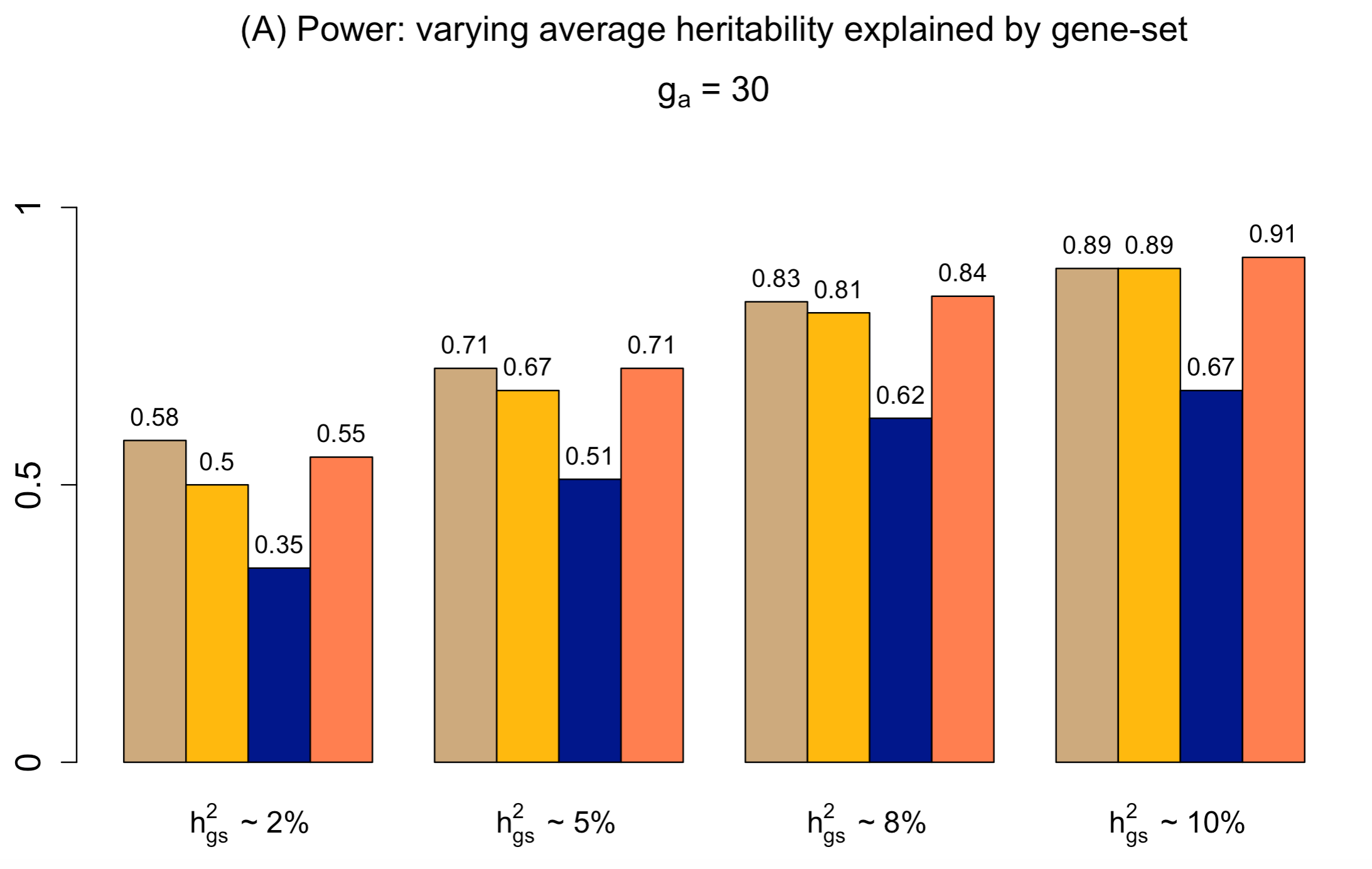

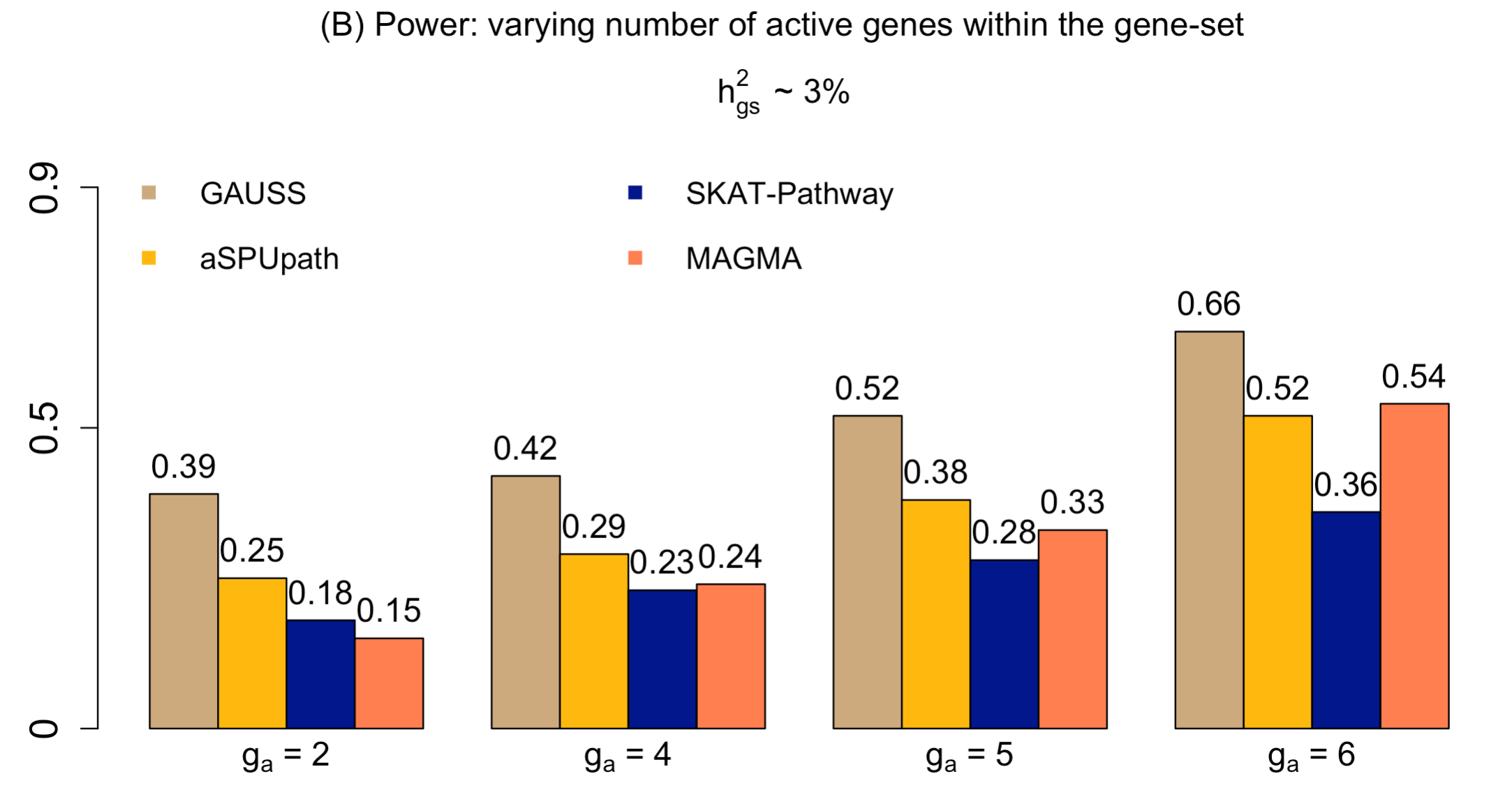

**Supplementary Figure 2**: **Power of GAUSS for GO: 0006955**. Estimated power of GAUSS using GO: 0006955 gene-set (1100 genes), compared with that of aSPUpath, SKAT-Pathway and MAGMA under different average heritability explained (*h^2^*) and different number of active genes (*g_a_*). (A) Power of GAUSS when 30 genes are active (*g_a_* = 30) and the variants with different average heritability (*h_gs_^2^*) explained by the gene-set. (B) Power of GAUSS with different number of active genes and the gene-set has an average heritability *h_gs_^2^* of 5%. The proportion of causal variants in an active gene (See Simulation Model) was set to 30%.

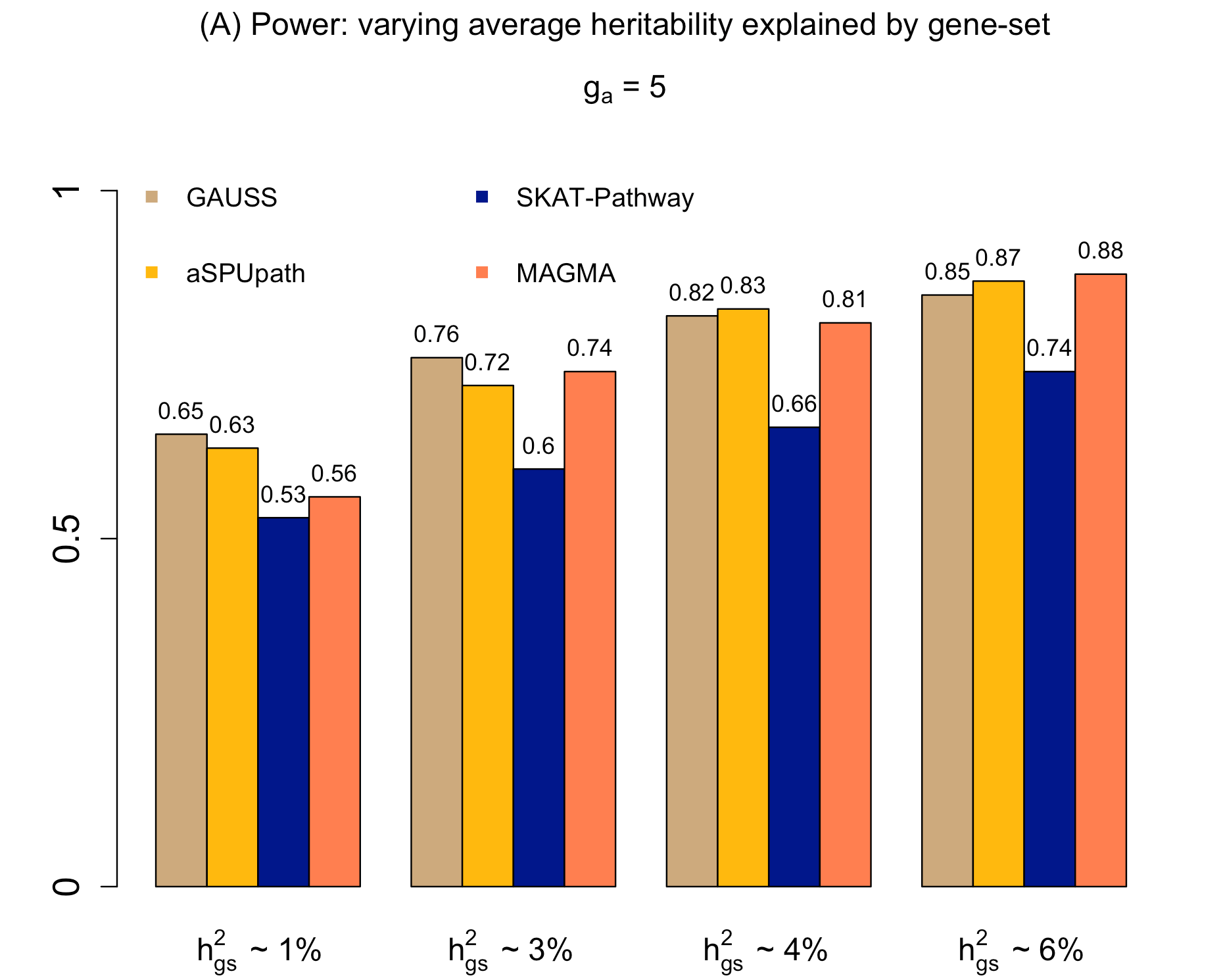

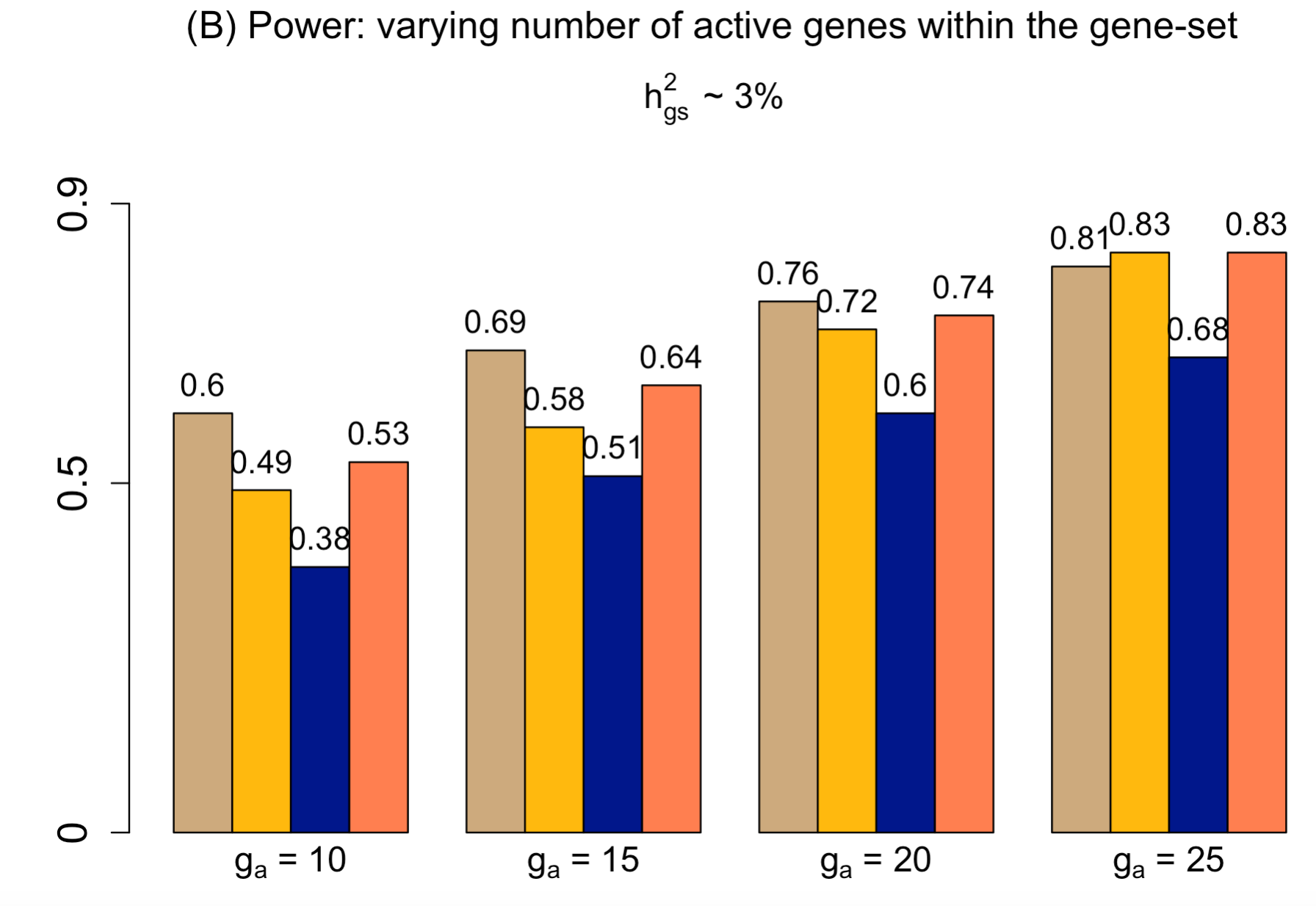

**Supplementary Figure 3**: **Power of GAUSS for GO:0002016**. Estimated power of GAUSS using GO: 0002016 gene-set (11 genes), compared with that of aSPUpath, SKAT-Pathway and MAGMA under different average heritability explained (*h^2^*) and different number of active genes (*g_a_*). (A) Power of GAUSS when 5 genes are active (*g_a_* = 5) and the variants with different average heritability (*h_gs_^2^*) explained by the gene-set. (B) Power of GAUSS with different number of active genes and the gene-set has an average heritability *h_gs_^2^* of 5%. The proportion of causal variants in an active gene (See Simulation Model) was set to 30%.

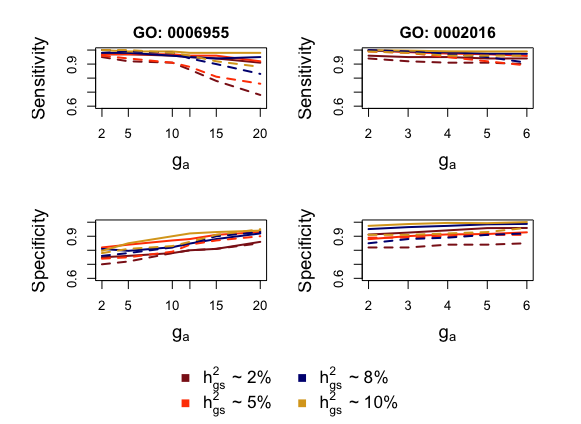

GAUSS

Significant Genes

**Supplementary Figure 4.** **Sensitivity and Specificity of GAUSS** **for GO: 0006955 and GO: 0002016**. Sensitivity and Specificity of GAUSS for GO: 0006955 (1100 genes) and GO: 0002016 (11 genes) with different average heritability explained by the gene-set (*h^2^*) and different number of active genes (*g_a_*). The solid lines denote GAUSS and the dashed lines denote the method of selecting the significant genes. The proportion of causal variants in an active gene (See Simulation Model) was set to 30%.

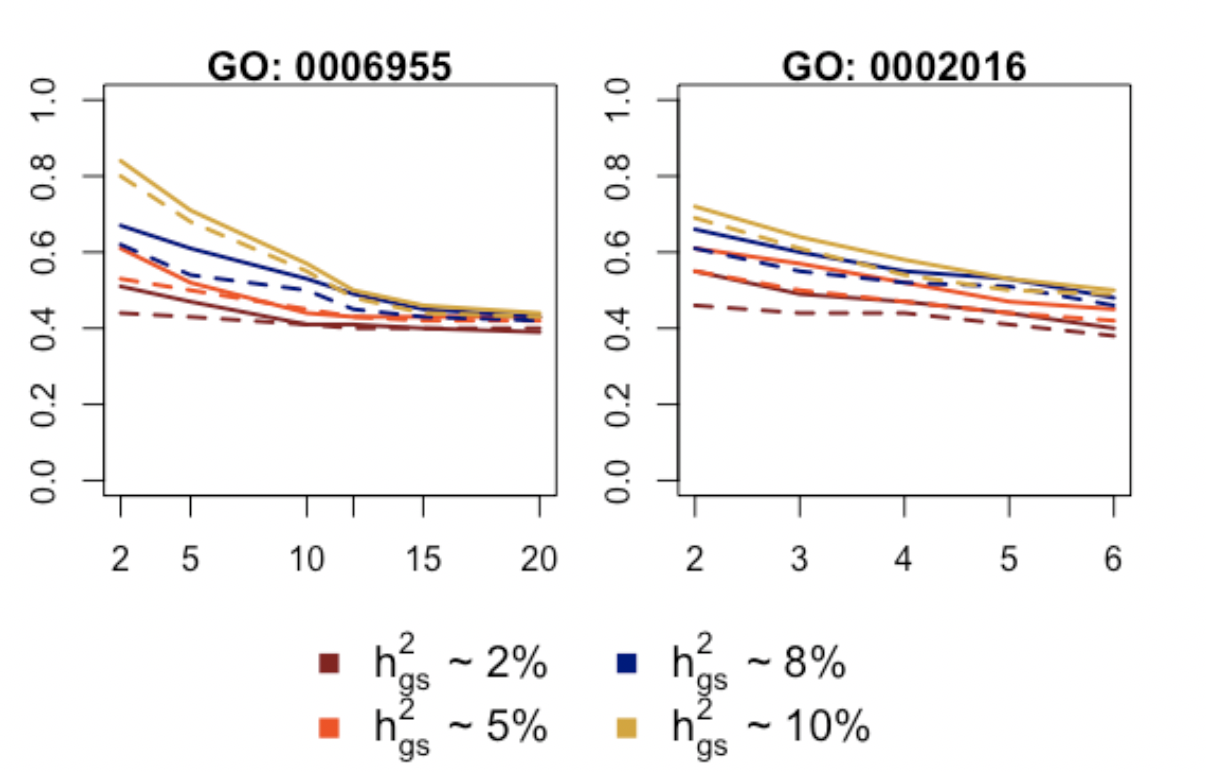

GAUSS

Significant Genes

**Supplementary Figure 5.** **Empirical power to identify correct non-null subset**. Estimate of the probability of identifying the exact set of non-null genes through CS genes, for different average heritability explained by the gene-set (*h_gs_^2^*) and different number of active genes (*g_a_*). The solid lines denote GAUSS and the dashed lines denote the method of selecting the significant genes.

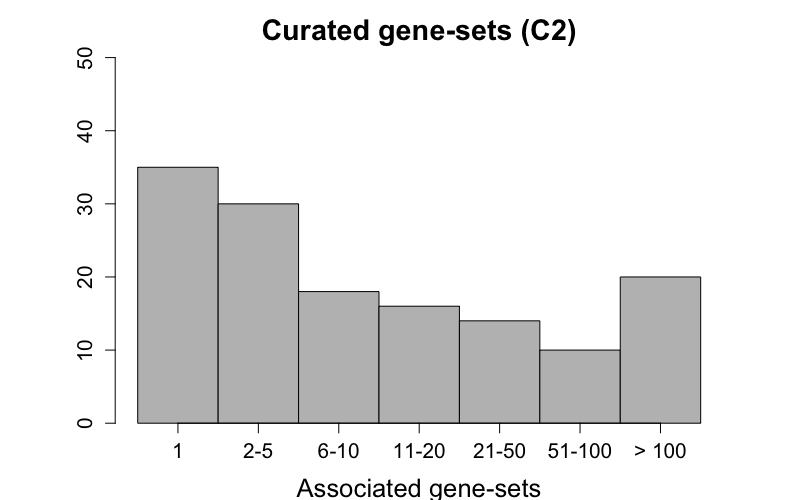

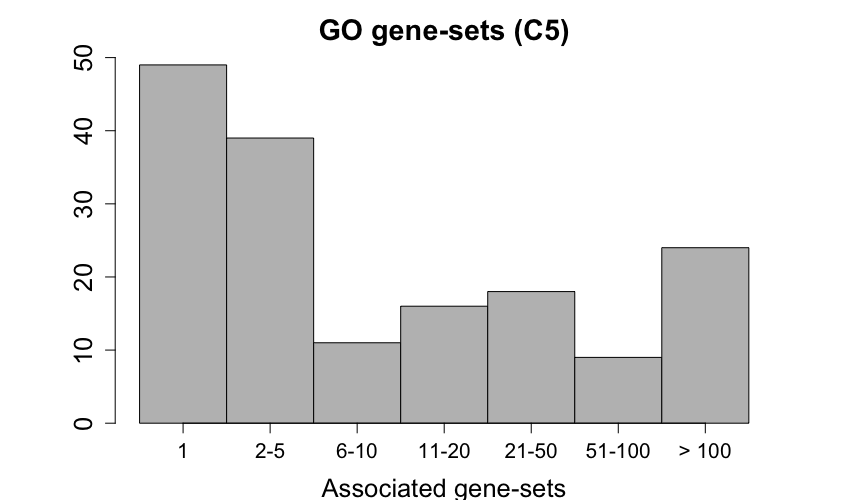

**Supplementary Figure 6**: Distribution of the number of significant gene-sets associated with phenotypes which had at least one significant gene-set in Curated gene-sets (C2) and GO gene-sets (C5).

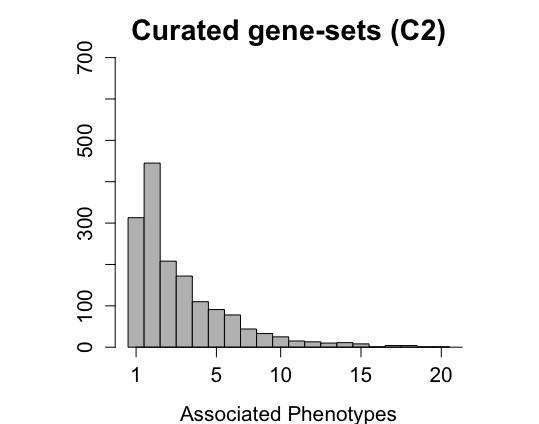

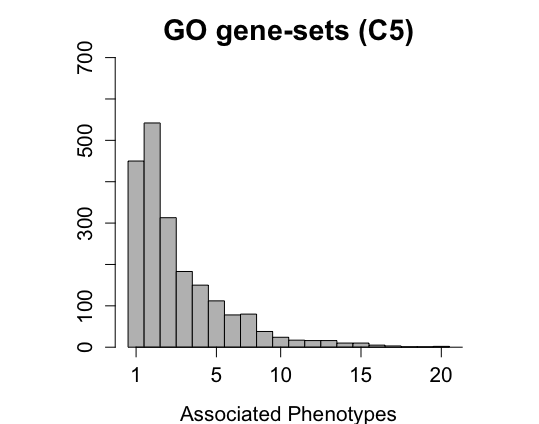

**Supplementary Figure 7**: Distribution of the number of significant phenotypes associated with gene-sets which had at least one significant phenotype.

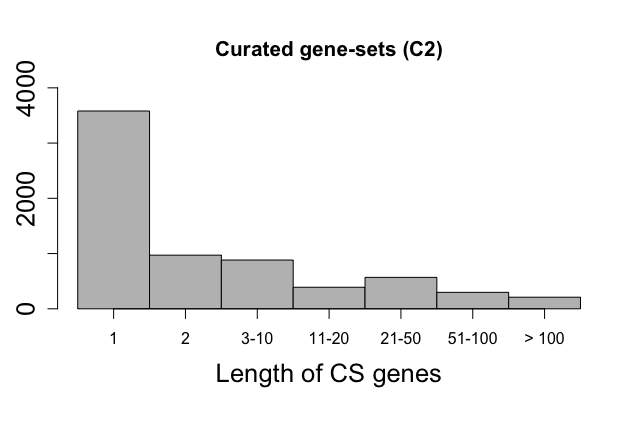

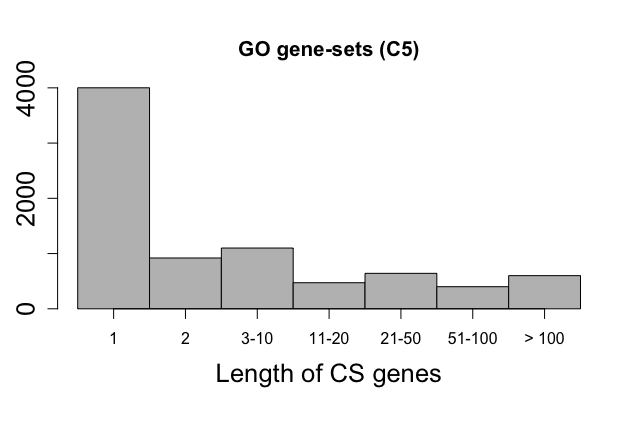

**Supplementary Figure 8**: Length of the core subset (CS) genes obtained by GAUSS for all the associations detected in UK-Biobank analysis

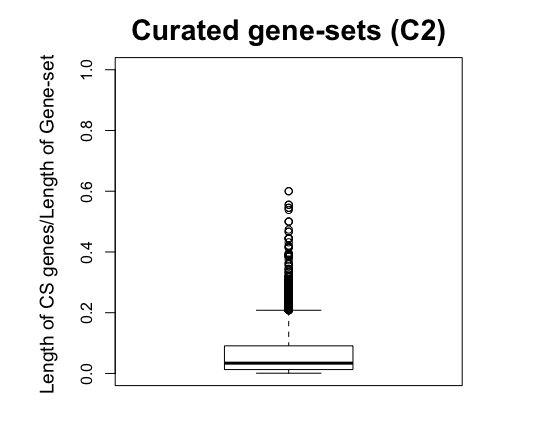

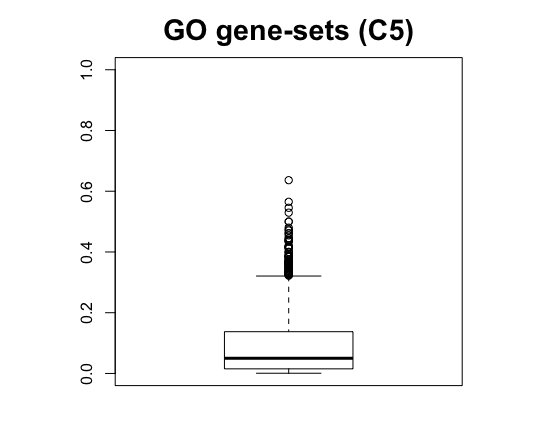

**Supplementary Figure 9**: Ratio of the length of the CS genes selected by GAUSS to the length of the gene-sets for Curated gene-sets (C2) and GO-gene-sets (C5). The figure only includes the significant associations.

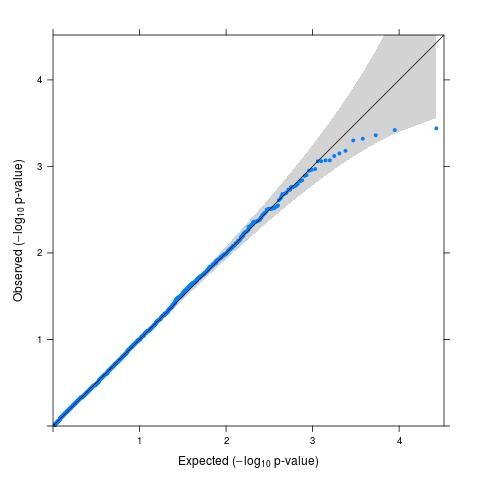

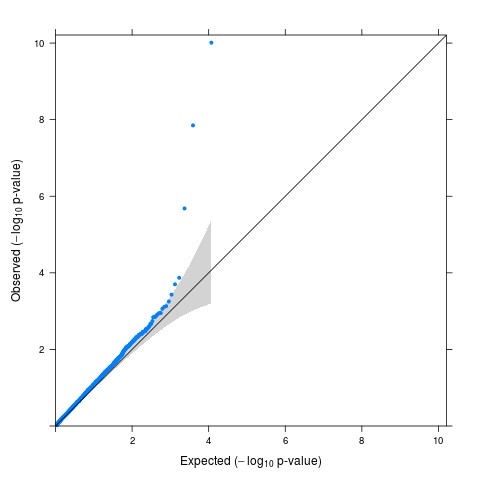

**Supplementary Figure 10**: QQ plots for gene-based p-values of E. Coli infection (EC; PheCode 041.4; Left Panel) and Gastritis and duodenitis (GD; PheCode 535; Right Panel)

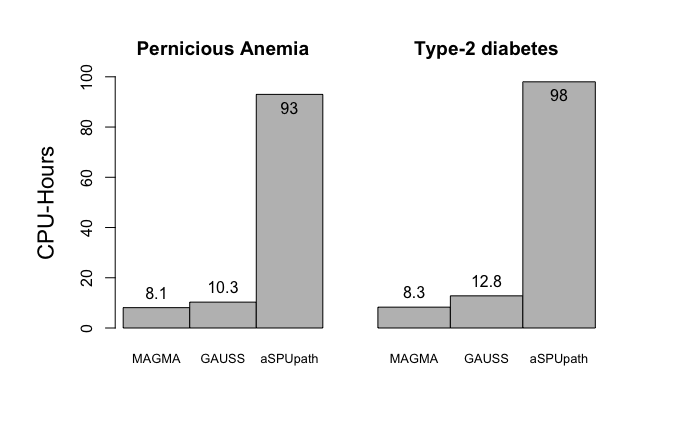

**Supplementary Figure 11**: Total run-time of GAUSS for Pernicious anemia and Type-2 diabetes in UK Biobank compared to that of MAGMA and aSPUpath. Total run-time is calculated as the clock-time taken starting from the input of summary statistic till the p-values for the 10*,*679 gene-sets are generated.

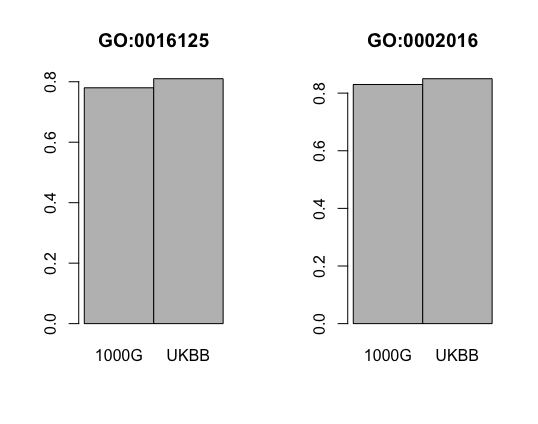

**Supplementary Figure 12**: Power of GAUSS using 1000-Genomes data (1000G) and UK Biobank data (UKBB) as reference panel. In this simulation, we first set the number of active genes to be 20 and 6 respectively in GO: 0016125 and GO: 0002016. The average h2 was set to be 4% and 30% of the variants within an active gene was set to have non-negative effect sizes. We estimated the LD from two different sources (1000-Genomes reference panel & UK Biobank reference panel) to construct the SKAT-Common-Rare tests. Then, we again estimated $\hat{V}_{H}$ from the same two reference panel two construct the GAUSS test. Here we compare the power of GAUSS corresponding to the two reference panels.
